# Molecular Basis of Maintaining Circannual Rhythm in the Skin of Cashmere Goat

**DOI:** 10.1101/2020.04.04.023044

**Authors:** Jianghong Wu, Ying li, Husile Gong, Dubala Wu, Chun li, Bin Liu, Lizhong Ding

## Abstract

The cashmere goat (*Capra hircus*) is famous for the fine quality cashmere wool. The cashmere is produced by secondary hair follicle that the growth shows seasonal rhythm. Thus, in this study, the skin of cashmere goat was selected as a model to illustrate the circannual rhythm of skin. The whole length skin transcriptome mixed from selected four months was obtained by PacBio single-molecule long-read sequencing (SMRT) technology. We generated 82,382 high quality non-redundant transcripts belonging to 193,310 genes, including 4,237 novel genes. Other 39 skin transcriptomes sampled from Dec. 2014 to Dec. 2015 were sequenced by Illumina Hi-Seq2500, we found 980 genes were differentially expressed. Of these genes, 403 seasonal rhythm genes (SRGs) were expressed and exhibited a seasonal pattern in skin. The results also showed that miRNAs were differentially expressed as the daylight length changed throughout a year. Some SRG genes related to the hormone secretion and eyes morphogenesis were enriched in skin. These genes gradually increased their expression level under short light, reached the peak near the summer solstice, and then began to decline. We found that the expression of Dio1 gene may be affected by the photoperiod that induces transformation from the inactive T4 to active thyroid hormone T3 in the skin and led to the difference between the skin circannual rhythm and the core circannual rhythm. Furthermore, the skin expressed eye morphogenesis-related genes and miRNAs, which suggested some cells in the skin could have the potential of light sensitivity. These results revealed that SRGs could regulate the downstream gene expression and physiological process in the skin to adapt to the season change.

## Introduction

To survive and reproduce, all living organisms must adapt to environmental variation. Thousands of genes of human blood have seasonal expression profiles, with inverted patterns observed between Northern and Southern hemispheres [1]. Thus, the gene expression of organisms would be regulated by the factors of the environment. Among these factors, the photoperiod is the most important clue for animals to adjusting the physiological state to the circadian and circannual rhythms that resulted from rotation and revolution of earth.

Although the eye is very important in the formation of rhythms, the skin is the largest tissue of the organism, which has the functions of defense, protection, and perception (such as the perception of seasonal changes). In the long evolution of living things, skin adapts to different habitats (water to land to sky, wet to dry, hot to cold) and evolves different types of skin. When multicellular organisms evolved to Annelida, they still didn’t have eyes. Before the appearance of special photosensitive organs, the skin had played a role in sensing the external environment (light). Light can directly affect the cell growth cycle [2]. Light can directly regulate the expression of a biological clock gene *PER* in cells [3]. Further studies have found that the phenomenon of the direct control of circadian clock by light without suprachiasmatic nucleus (SCN) is common in fish [4, 5]. Miki Tanioka et al. first found fluctuating biomolecular clocks in the skin of mice [6]. However, subsequent studies found that both rat dermal fibroblasts [7] and human keratinocytes cultured in vitro have the same rhythm as the peripheral biological clock, and the external environment (drugs, serum) can affect the biological clock of cells cultured in vitro[8–10]. The circadian clocks of individual cells cultured in vitro are controlled by themselves, and the circadian clocks can be transmitted to the progeny cells through the process of cell division and proliferation [11]. Different types of cells in the skin have different biological clock phases. The whole biological clock in the skin is formed by the coordination of various types of cell biological clock genes [12]. Therefore, the biological clock in the skin is very complex and has memory potential, which is easily affected by light.

The economic performance of livestock is mostly related to seasonal rhythm, especially the animals living in the temperate zone. Animals rely on environmental cues to judge the coming season, in order to synchronously change physiological conditions and phenotypic behavior along with environmental condition alterations, It is generally accepted that mammals perceive the light and darkness of the outside world through the retina and transmit the photoperiod messages to the hypothalamus through the content of melatonin in the pineal gland, which depends on light to control the physiological rhythm. So the biological clock in the body keeps the same rhythm with the environment. The circannual rhythms are essential for animal production. Therefore, the study on the mechanism of how livestock adjusts gene expression in different tissues to adapt the biological systems to seasonal changes by changing their physiological behavior. This study can provide a theoretical basis for animal husbandry production.

## Results

### Skin transcriptome of cashmere goat skin

To obtain a representative whole length transcriptome of skin, we collected body side skin and ear skin from three Inner Mongolia Cashmere Goats at four time points (Jan, Apr, Jul, Oct) and employed PacBio single-molecule long-read sequencing (SMRT) technology. 591,585 circular consensus (CCS) reads were obtained. Of the total reads, 466,058 (78.78%) full length reads non-chimeric (FLNC) sequences were identified. After Iterative Clustering, 149,604 consensus isoforms were polished and 137,211 high quality FLNC transcripts were classified, with the criteria post-correction accuracy above 99%. The high-quality Illumina sequencing was used to improve the quality of long reads. Finally, 82,382 high quality and non-redundant transcripts were obtained, which belong to 18,919 genes, including 4,237 novel genes.

**Figure 1.**
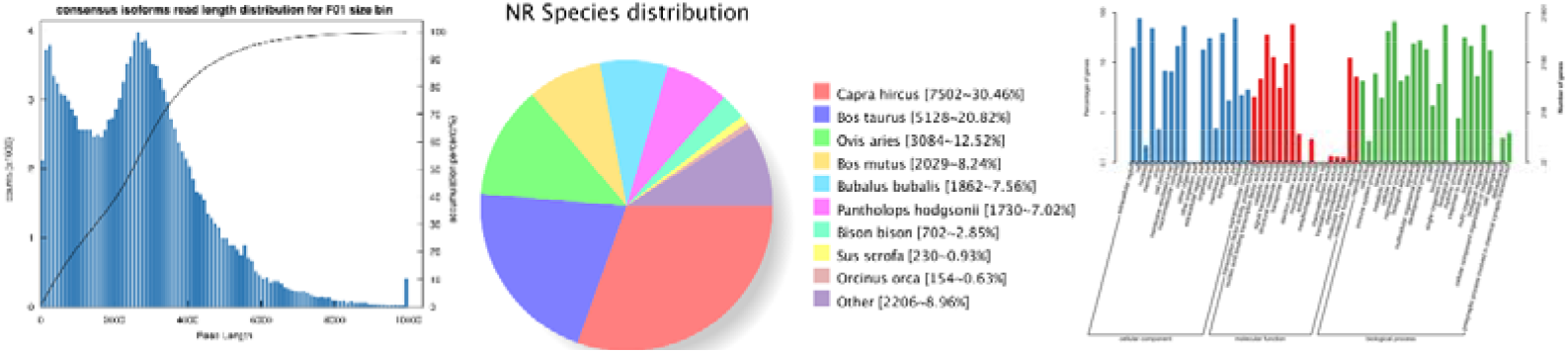
Summary of PacBio RS II SMRT sequencing of *cashmere goat skin* transcripts. **(a)** consensus isoforms read length distribution, **(b)** the species distribution of gene annotation based the NR database, **(c)** GO annotation, where blue represents a cellular component, red represents a molecular function, and the green represents a biological process.

To reveal Annual Rhythm under natural conditions, we chose Cashmere goats, strict seasonal molt livestock as the study species. After the reads filtering, clean data were obtained. To calculate the expression level of skin genes, Salmon [13] software was utilized based on the whole length transcriptome reference sequence. Finally, 980 genes have significantly differential expression in goat skin among the different months. The sample correlation matrix showed that the transcriptomes of goat skin were divided into long daylight (April, May, Jun, July, August) and short daylight group (September, October, November, December, January, February, March) (Figure 2). This result suggested that these differential expressed genes (DEGs) could be expressed in the skin under the certain rhythm.

**Figure 2.**
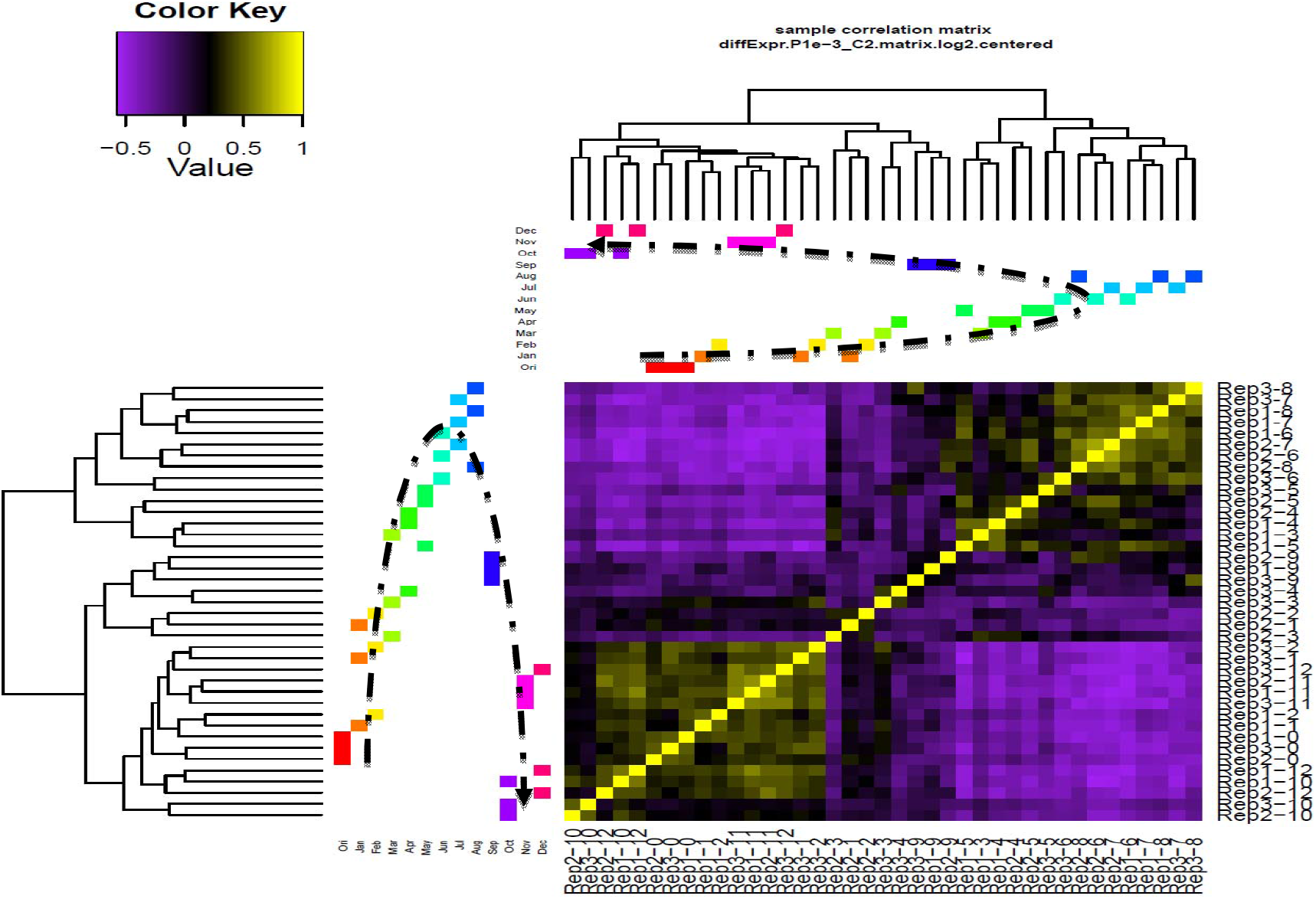
The Sample Correlation Matrix Based on the Skin Transcriptomes. Those colorful blocks represent the biological replicates at different time point. The curve shows that the skin transcriptome profile dynamic change with the seasonal variation.

To understand the dynamics of chromosome expression, we analyzed the distribution of DEGs on each chromosome and calculated the density of DEGs. The results showed that the genes on the Chr 1 and Chr 19 were significantly activated. Furthermore, we found the KP and KAP tandem region on those two chromosomes (Figure 3).

**Figure 3.**
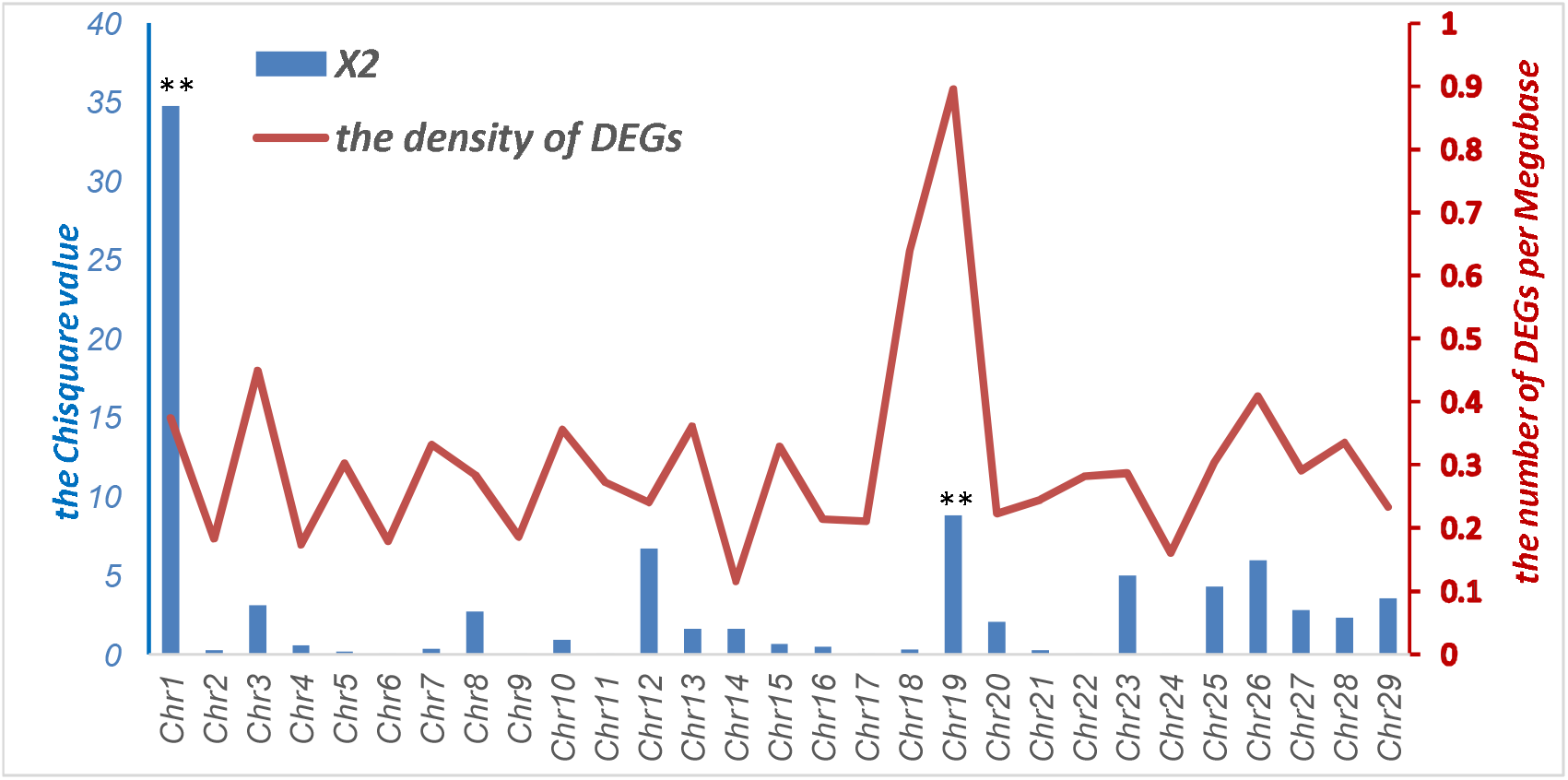
the distribution of DEGs on the chromosome.

### Annual Rhythm Expressed Genes in Goat Skin

To analyze the rhythm of DEGs of goat skin, MetaCycle [14] was utilized. We tried from 4 months to one year as a length of interesting rhythms. Finally, 403 genes of those DEGs expressed in the skin as circannual rhythm (Figure 4), which are divided into two groups long daylight genes and short daylight genes.

**Figure 4.**
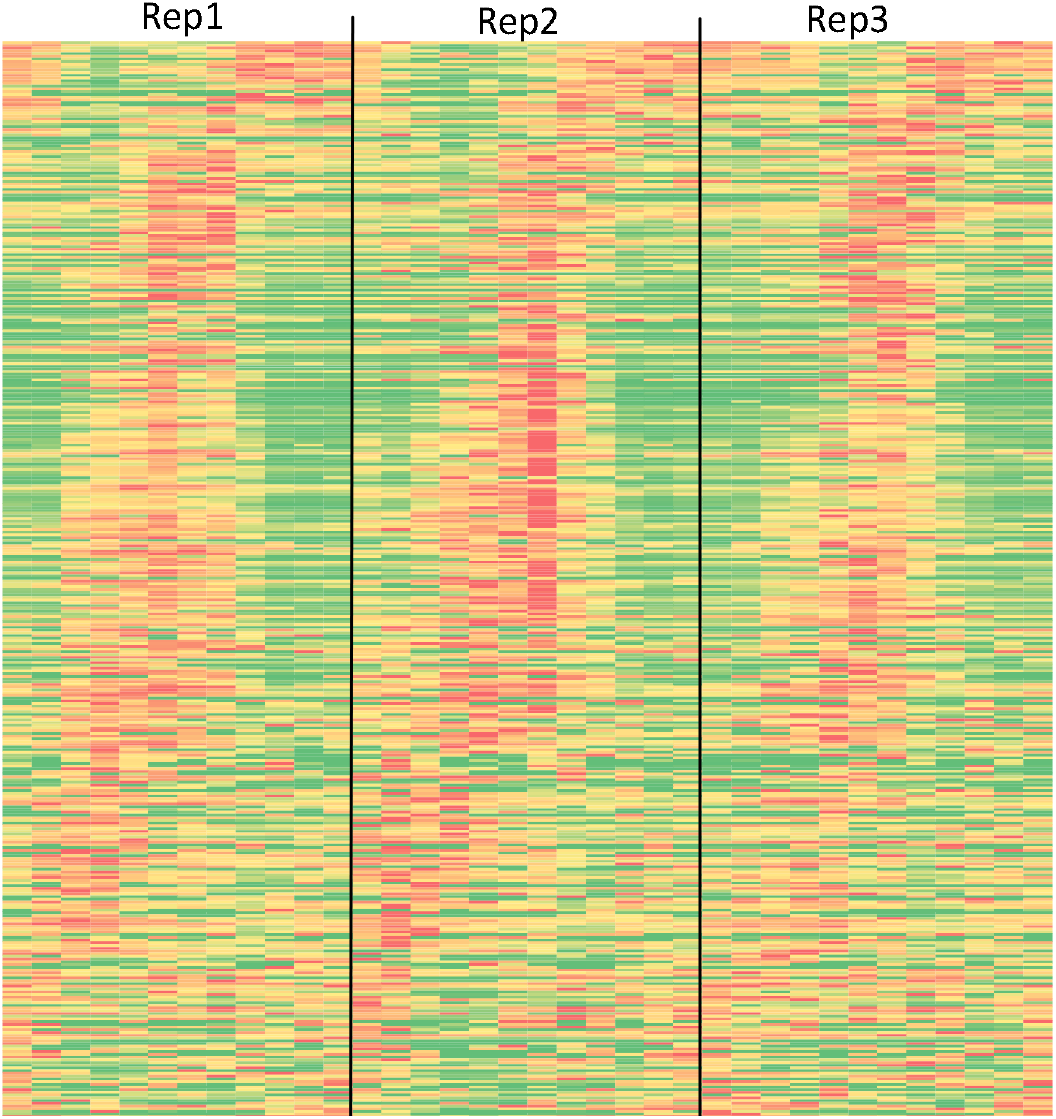
The Heat Map of The Seasonal Rhythm Genes in Cashmere Goat Skin. Rep1, Rep2 and Rep3 are the three replicates of experimental goat.

We identified 90 transcription factors (TFs) and 76 TF cofactors (Supplement file 1) in these seasonal rhythm genes (SRGs) of goat skin from the AnimalTFDB [15]. Of these SRGs, 18 ncRNAs were highly expressed in skin. And results also showed that there are 28 genes related to the secretoglobin family and S100 family with high expression (Table 1). Furthermore, to infer the potential upstream regulators of SRGs, we investigated the transcript binding site on the 5’ UTR of SRGs. Many of the ZNF motif and the KLF motif are located in the FCR of SRGs (Supplement file 1). KLF family members are related to the pathways of BMAL1-CLOCK and NPAS2, which could activate circadian gene expression and Adipogenesis.

**Table 1.** The genes related with secretoglobin family and the S100 family.

| RefSeq Accession Number | Gene symbols | Gene description | Maximum expression value (RPKM) |
| --- | --- | --- | --- |
| XM_005692259.2 | LOC102169125 | major allergen I polypeptide chain 2 | 45989.907 |
| XM_018062299.1 | LOC106503120 | major allergen I polypeptide chain 1 | 45277.958 |
| XM_005692256.2 | LOC102191166 | major allergen I polypeptide chain 1-like | 40423.136 |
| XM_013971277.2 | LOC102191453 | secretoglobulin family 2B member 2-like | 32363.733 |
| XM_018062302.1 | LOC108637981 | major allergen I polypeptide chain 1-like | 31126.557 |
| XM_018044050.1 | LOC102188300 | odorant-binding protein | 11397.749 |
| XM_005692260.3 | LOC102169411 | major allergen I polypeptide chain 1-like | 10994.02 |
| XM_018044026.1 | LOC102186944 | allergen Bos d 2 | 9852.28 |
| XM_005692261.2 | LOC102169695 | major allergen I polypeptide chain 2 | 9054.857 |
| XM_005677509.2 | LOC108633164 | protein S100-A7-like | 7418.243 |
| XM_005701239.2 | LOC102189201 | allergen Bos d 2 | 6038.088 |
| XM_005677510.2 | LOC108633164 | protein S100-A7-like | 3554.848 |
| XM_018046586.1 | S100A7A | protein S100-A7A | 2872.84 |
| XR_001919671.1 | LOC102180344 |  | 2441.43 |
| XM_005692255.2 | LOC102190903 | androgen-binding protein homolog | 2432.44 |
| XM_018046587.1 | S100A7A | protein S100-A7A | 2122.853 |
| XM_018044027.1 | LOC102186944 | allergen Bos d 2 | 2112.612 |
| XM_005677513.3 | S100A8 | protein S100-A8 | 1697.389 |
| XR_311282.2 | LOC102187389 |  | 1392.066 |
| XM_018046133.1 | LOC102169149 | protein S100-A12 | 1185.225 |
| XM_013977249.2 | S100A9 | protein S100-A9 | 566.149 |
| NM_001285548.1 | LTF | lactotransferrin precursor | 312.126 |
| XM_018065147.1 | LOC102182869 | primary amine oxidase, lung isozyme-like isoform X1 | 299.292 |
| XR_310700.1 | LOC102191729 |  | 270.069 |
| XM_018062304.1 | LOC102168852 | major allergen I polypeptide chain 1-like | 247.89 |
| XM_018046132.1 | S100A9 | protein S100-A9 | 81.69 |
| XM_005679934.3 | KRT4 | keratin, type II cytoskeletal 4 | 64.26 |
| XM_005693167.3 | LOC102179878 | WAP four-disulfide core domain protein 18 | 62.968 |

### The molecular basis of hair growth

To character the hair cycle on the molecular level, we chosen keratin protein (KP) genes and keratin-associated protein (KAP) genes from the DEGs, which are the majority of cashmere composition. In this study, of those DEGs, 55 genes of KP and KAP were highly expressed in goat skin during the changing of the season. 37 transcripts reached the peak of expression level at August and highly expressed during June to September, which was earlier than the hair follicle activity (Figure 5). The majority of these genes are located on the Chr1:141.4M-144.5M and Chr19:41.0M-41.3M, according to the chromosomal distribution of DEGs.

**Figure 5.**
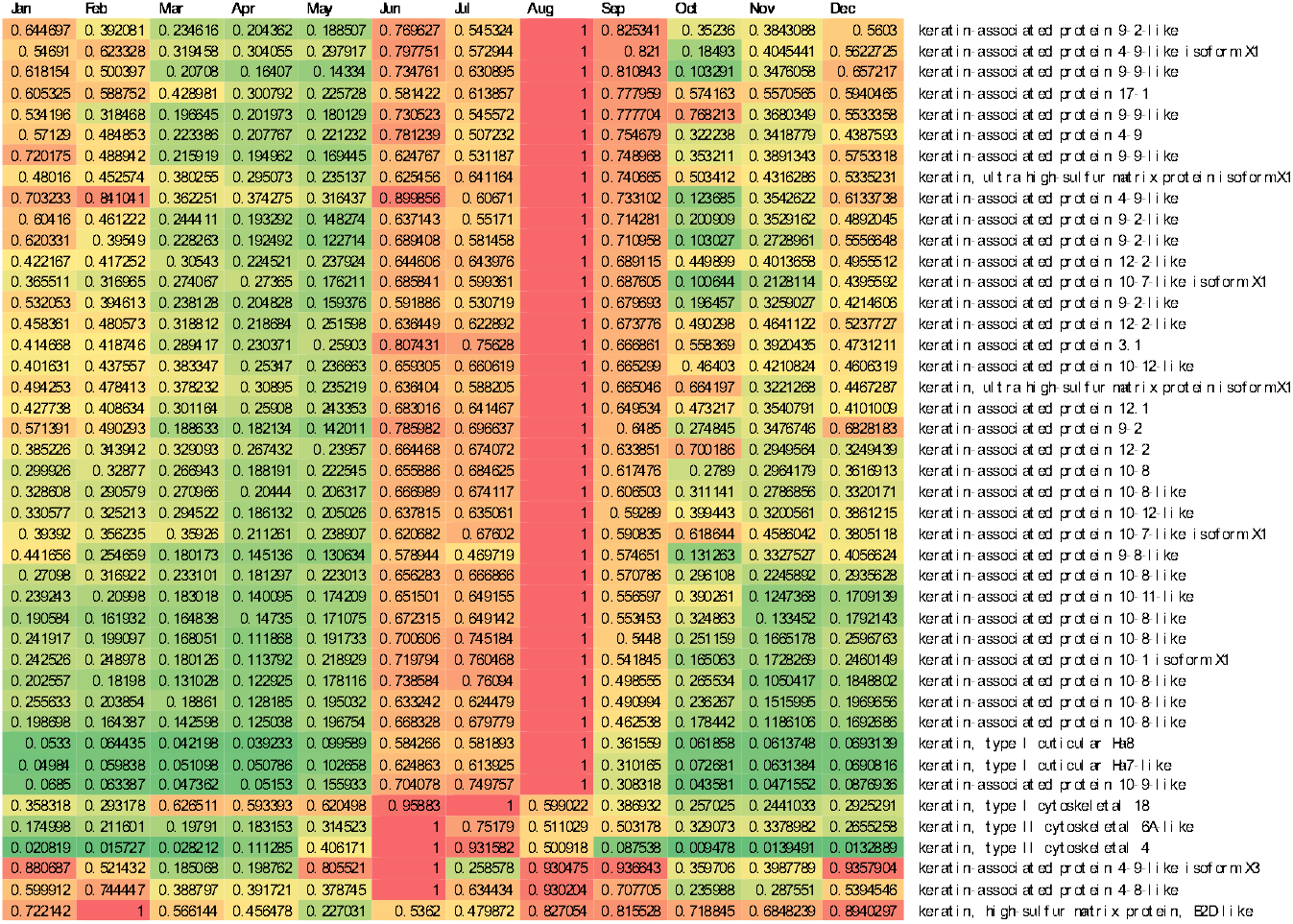
The Expression Matrix of the Keratin Protein Genes and Keratin-associated Protein Genes in Skin at One Year.

#### The function enrichment of seasonal rhythm genes

The function of seasonal rhythm genes in the skin was enriched by the ClueGO, and 18 clusters GO terms were classified. Most of the GO terms are involved in the biological process of organ growth and development, signal transduction, and response stimulate from activity components (Figure 6A). Of these GO clusters, eye morphogenesis and camera-type eye morphogenesis were enrichment in cashmere goat skin. Therefore, we interest in whether the skin also has the sensor for light. To deeply analyze the expression pattern of eye morphogenesis genes, we drew the expression value of those genes related to the processes. The majority of these genes were highly expressed in the skin during in long daylight of year (Figure 6B).

**Figure 6.**
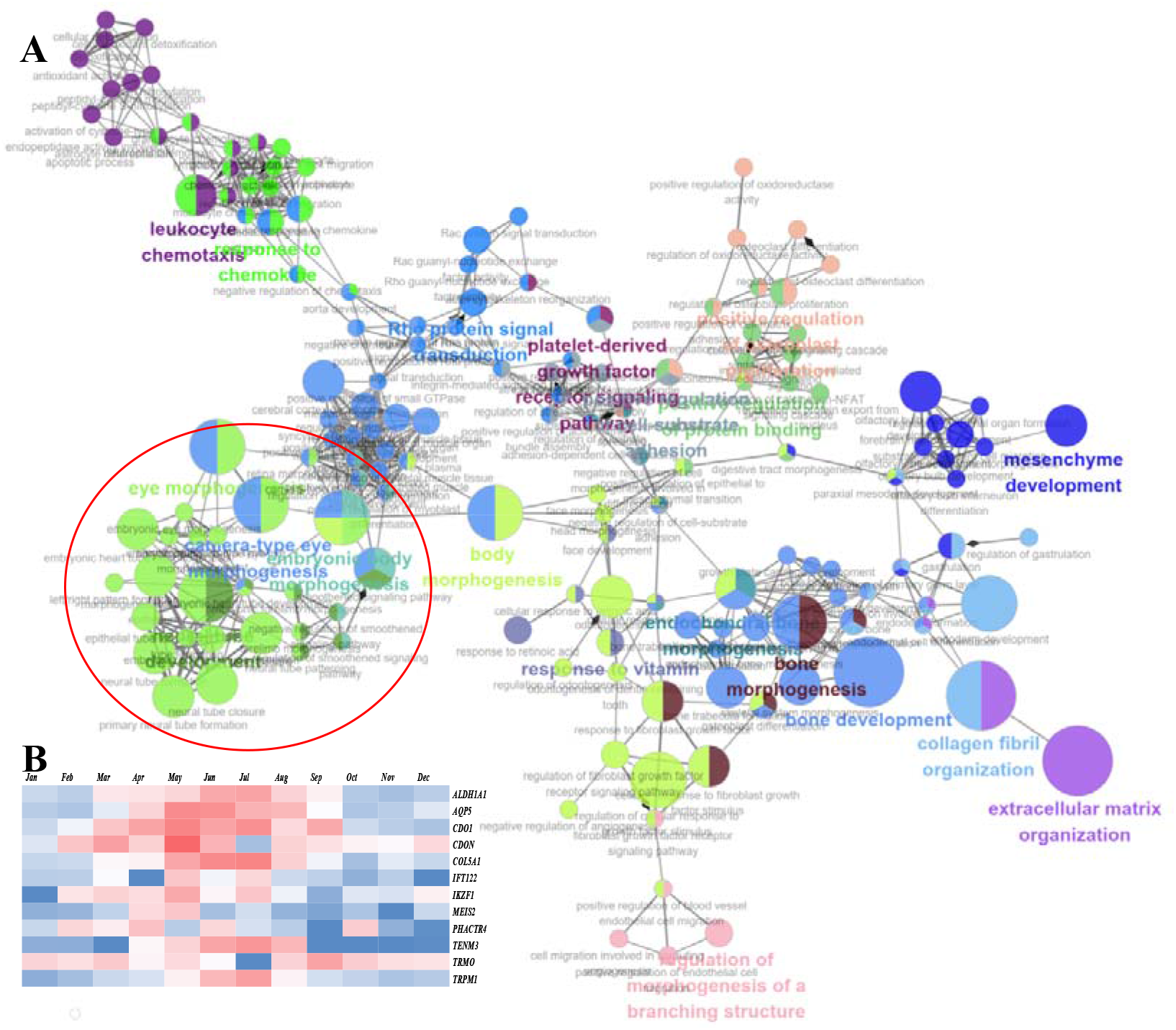
The Function Enrichment and Expression Pattern. **A,** the Function Enrichment for Annual Rhythm Expressed Genes based on the GO (biological process), the red circle points out those biological process related with the development of eye. **B**, the expression pattern of eye morphogenesis genes.

The thyroid hormone signaling pathway and thyroid hormone synthesis were enriched by ClueGO (Figure 7A). Three types deiodinase, DIO1, DIO2, and DIO3, were expressed in the skin of cashmere goats (Figure 7B), which was essential to the regulation of thyroid hormone activity. To deeply analyze the expression pattern of thyroid hormone related genes, we drew the expression value of those genes related to the processes. The expressed trend of these genes showed a certain rhythmic pattern in the skin during the whole year (Figure 8).

**Figure 7.**
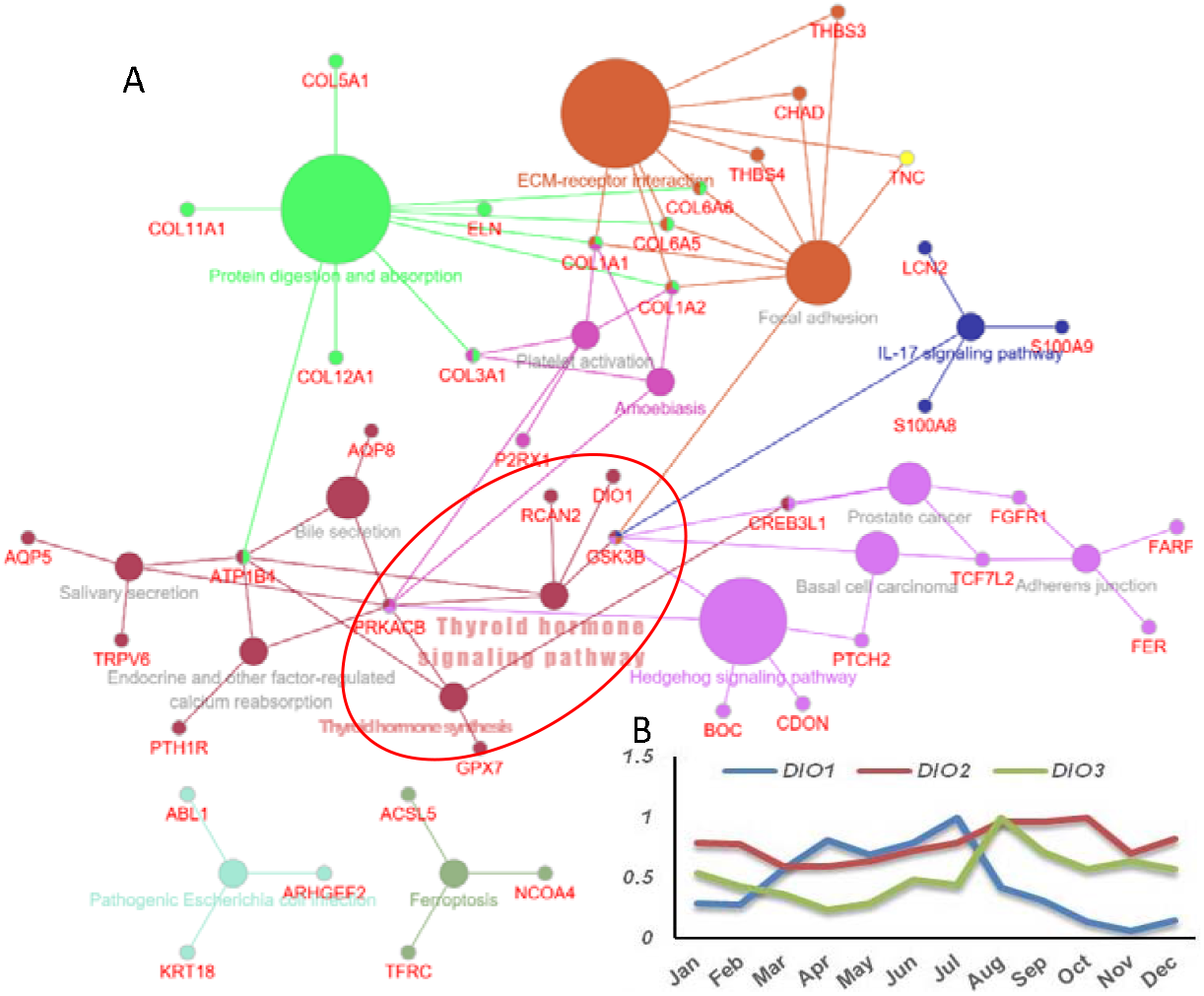
The Function Enrichment for Annual Rhythm Expressed Genes and the Expression of the Member of Deiodinase. **A**, the function enrichment for annual rhythm expressed genes based on the KEGG, the red circle points out thyroid hormone pathway and genes. **B**, the expression of the member of deiodinase in goat skin.

**Figure 8.**
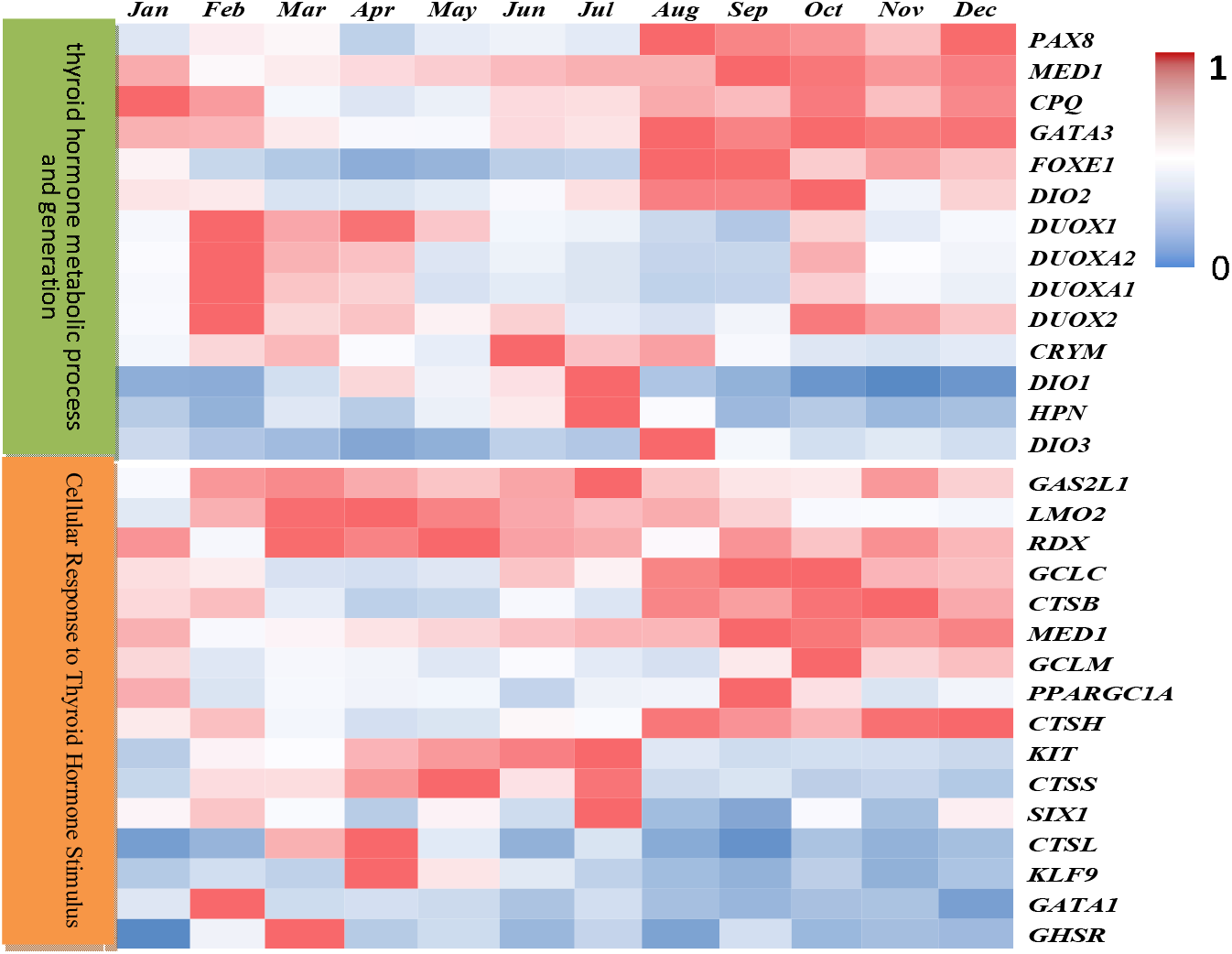
The Expression Pattern of Thyroid Hormone Associate Genes.

#### The differential expressed miRNA in goat skin during seasonal variation

To investigate whether the miRNA would variate during seasonal changed, we also used NGS to sequence the miRNA in cashmere goat skin. 1973 mature miRNAs were predicted based on the miRDeep2 software [16], which contains 436 known goat mature miRNAs and 1537 novel miRNAs. 35 miRNAs were differentially expressed between winter and summer, which was the biggest difference among four seasons (Table 2). Of these differential expressed miRNAs (DEMs), 33 miRNAs were up and 2 were down-regulated in winter, compared to summer (Supplement file 2). The chi-miR-206 was highly expressed in winter compared with the summer, which indicated that miR-206 is also an important regulator of the circannual rhythm excepting an important regulator of the circadian clock [17]. Chi-miR-211 and chi-miR-144 are expressed lower in winter compared with the summer.

**Table 2.** The number of DEMs among different seasons

|  | Spring | Summer | Fall | Winter |
| --- | --- | --- | --- | --- |
| Spring | 0 | 0 | 0 | 0 |
| Summer | 0 | 0 | 2 | 35 |
| Fall | 0 | 2 | 0 | 2 |
| Winter | 0 | 35 | 2 | 0 |

The expression trend of DEMs was opposite to the SRGs in skin during the seasonal variation. To integrate the miRNA and SRGs, we utilized the Mirnada to detect the potential binding sites. Of those SRGs, 151 genes have the interaction with 35 miRNAs based on the Mirnada. Then, the interaction networks were visualized in Cytoscape, which constructed from 186 genes (nodes) and 998 interactions (edges) (Figure 9).

**Figure 9.**
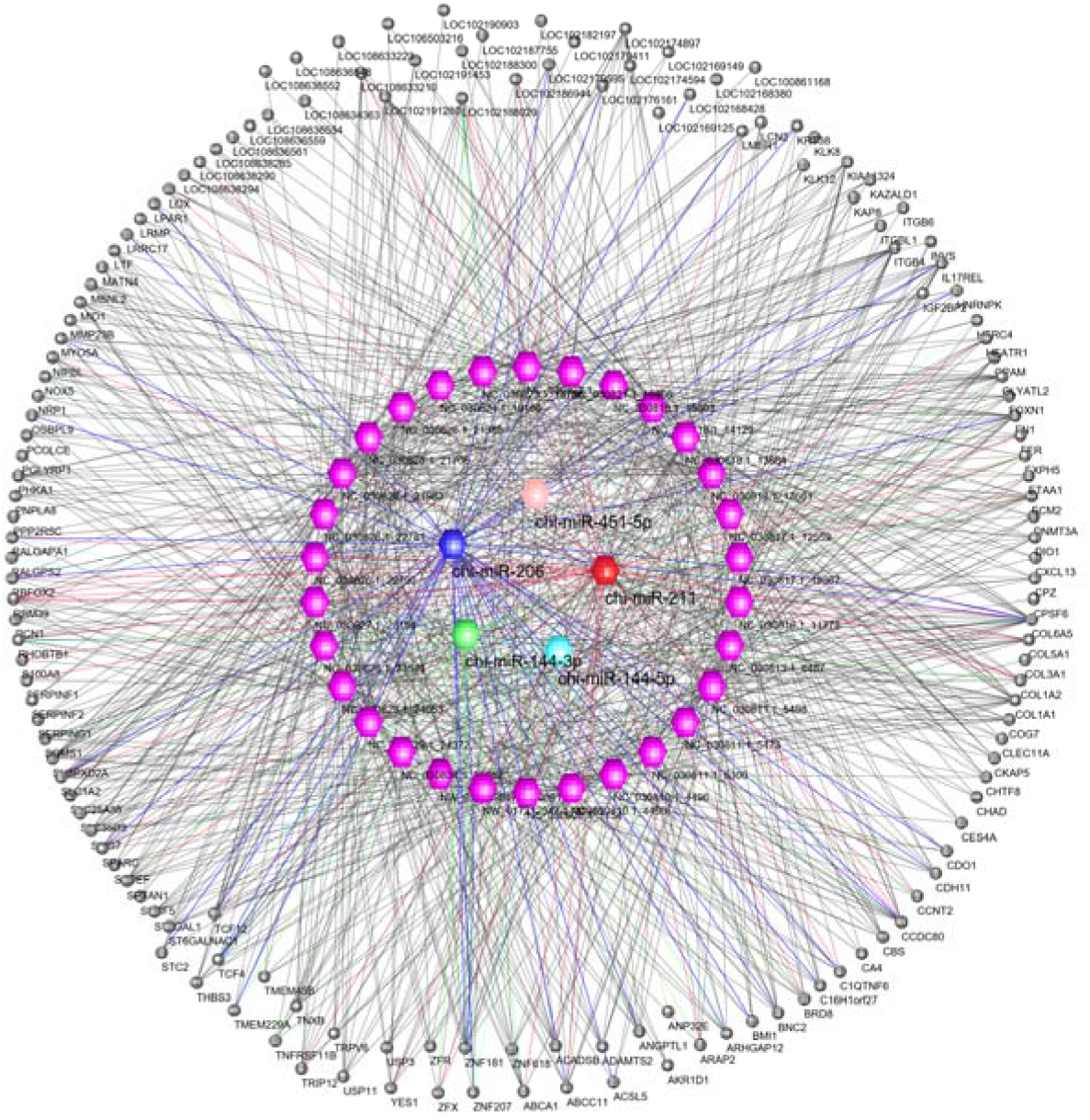
The Interaction Networks Between Seasonal Rhythmal Genes and Differential Expressed miRNAs During Seasonal Variation. Inner cycle for miRNAs and outer cycle for SRGs. The purple miRNAs were novel miRNAs based on the miRDeep2.

## Discussion

In this study, 403 seasonal rhythm genes were obtained by analyzing the dynamic transcriptome of cashmere goat skin. However, the classic circadian clock genes (ClOCK, BMAL1, PER, TIM, CRY, CSNK) [18] did not show significant seasonal changes in cashmere goat skin. The results suggested that there might be differences in gene sets controlling seasonal rhythm and circadian rhythm. However, many KLF motifs are located in the FCR of SRGs, which indicated that the regulation of the circannual rhythm in skin could depends on the circadian clock.

Through the functional classification of 403 seasonal rhythm genes (SRGs), it is found that these genes are involved in the synthesis of various hormones (thyroid hormones), sensory organ development (eyes), keratinocyte differentiation (such as epidermal differentiation complex gene cluster gene), and other functions. It is generally accepted that seasonal rhythms consist of complex transcription and translation feedback pathways, which activate specific transcription factors through thyroid hormones. The results showed that the coding genes of type I deiodinase (DIO1) was expressed in the skin of cashmere goats in the seasonal rhythm. The expression of DIO1 gene in cashmere goat skin increased gradually under short light but decreased gradually under long light. In addition to the ability to deionize T4 and transform it into T3, DIO1 can also remove the excessive thyroid hormone T4 in the tissue and protect the body from the damage caused by hyperthyroidism [19, 20]. The skin and hair follicle express almost all thyroxine receptor proteins [21–23], which shows that the skin can make precise regulation on seasonal thyroid hormones to adapt to seasonal changes and regulate the expression of transcripts in cells to achieve physiological behavior feedback of organisms. The expression of Dio1 in cashmere goat skin increased gradually under short light, reached the peak before and after the summer solstice, and then began to decline, which was equivalent to increasing the concentration of T3 in cashmere goat skin in autumn and winter. With the increase of T3 concentration in autumn and winter, the activity of secondary hair follicle stem cells increased, and then the growth of villi started.

CHGA gene belongs to the short light labeling of thyrotropin cells, and the long light thyrotropin cells labelled by the EYA3 gene maintain the annual biological rhythm of pituitary nodule through the transformation of two-phase state [24]. In this study, CHGA was expressed in the skin of cashmere goats in the seasonal rhythm keeping step with the Dio1 gene. Although the expression of CHGA in the skin of cashmere goat is not consistent with that in the pituitary nodule, the regulation of seasonal rhythm in the skin may be different from that in the brain. Of the SRGs, those genes related to the eye morphogenesis were enriched, such as AQP5 and TRPM1. Both of them are highly expressed during long daylight. AQPs plays important roles in all ocular tissues and involved in the eyes morphogenesis [25]. Mutations in TRPM1 (Transient Receptor Potential Cation Channel Subfamily M Member 1) could induce complete congenital stationary night blindness [26]. UVB could affect the activity of the p53◻LTRPM1/miR◻L211◻LMMP9 axis in human melanocyte and induce melanocyte migration [27]. In this study, we also found that chi-miR-211 was highly expressed in cashmere goat skin in summer. MiR-211 is essential for adult cone photoreceptor maintenance [28]. These results suggested that some cells in the skin could have the potential of photosensitive, which could maintain the specific rhythm of the peripheral organs based on the environment factor (photoperiod).

The results also showed that the epidermal differentiation complex (EDC) genes’ expression gradually increases under short light, reaches the peak at about the summer solstice, and then begins to decline. The main function of EDC is to drive keratinocyte differentiation and promote epidermal and hair follicle differentiation, which may be the reason why cashmere goats began to grow cashmere at the end of June and the beginning of July. Among them, S100A7, S100A8, S100A9 and S100A12 genes are all innate immune regulatory elements [29] in humans, indicating that skin plays an important role in organism protection and environmental perception. The level of S100A7 expressed in human normal skin is very low, and only under pathological conditions (such as melanoma) will have the expression increase [15, 30–32]. This may suggest that S100 gene family members or EDC gene cluster members of cashmere goats have different biological functions from humans and mice. The S100A7 of ungulates has accelerated evolution [33], thus S100A7 gene of cashmere goat may have evolved different functions from human and mouse. Furthermore, the S100A7 gene can be induced and expressed by ultraviolet light in vitro and in vivo, with a dose-dependent effect [34]. Ultraviolet radiation and retinoic acid both can stimulate normal keratinocytes to secrete S100A7 protein, and finally inhibit the differentiation of keratinocytes [35].

The expression pattern of S100A7 in the skin of cashmere goats was highly correlated with the changing trend of UV intensity in the growing place [33]. It has also been reported that S100A7 can regulate β-catenin signaling pathway [36], while Wnt signal cannot induce the expression of S100 family protein [37], which is single directional. Therefore, UV may induce the differentiation and development of skin and hair follicle by affecting the expression of specific genes in the skin of cashmere goat, and then promote the seasonal growth of villi. However, the accumulation of β-Catenin is very important for the activation and proliferation of hair follicle stem cells. When the thyroid hormone receptor (TR) is knocked out, the proliferation of hair and epidermis in mice is limited [38], and the expression of several miRNAs related to skin homeostasis is significantly reduced [39]. The expression of S100A8 and S100A9 in the skin of knockout mice treated with retinoic acid increased significantly [40], indicating that TRs receptor related pathways may be related to the regulation of S100 expression. Further, it was found that T3 could increase S100 expression in Schwann cells [41], which indicated that T3 could regulate the epidermal differentiation complex genes such as S100, and then regulate the activity of Wnt and other hair follicle development related signal pathways.

**Figure 10.**
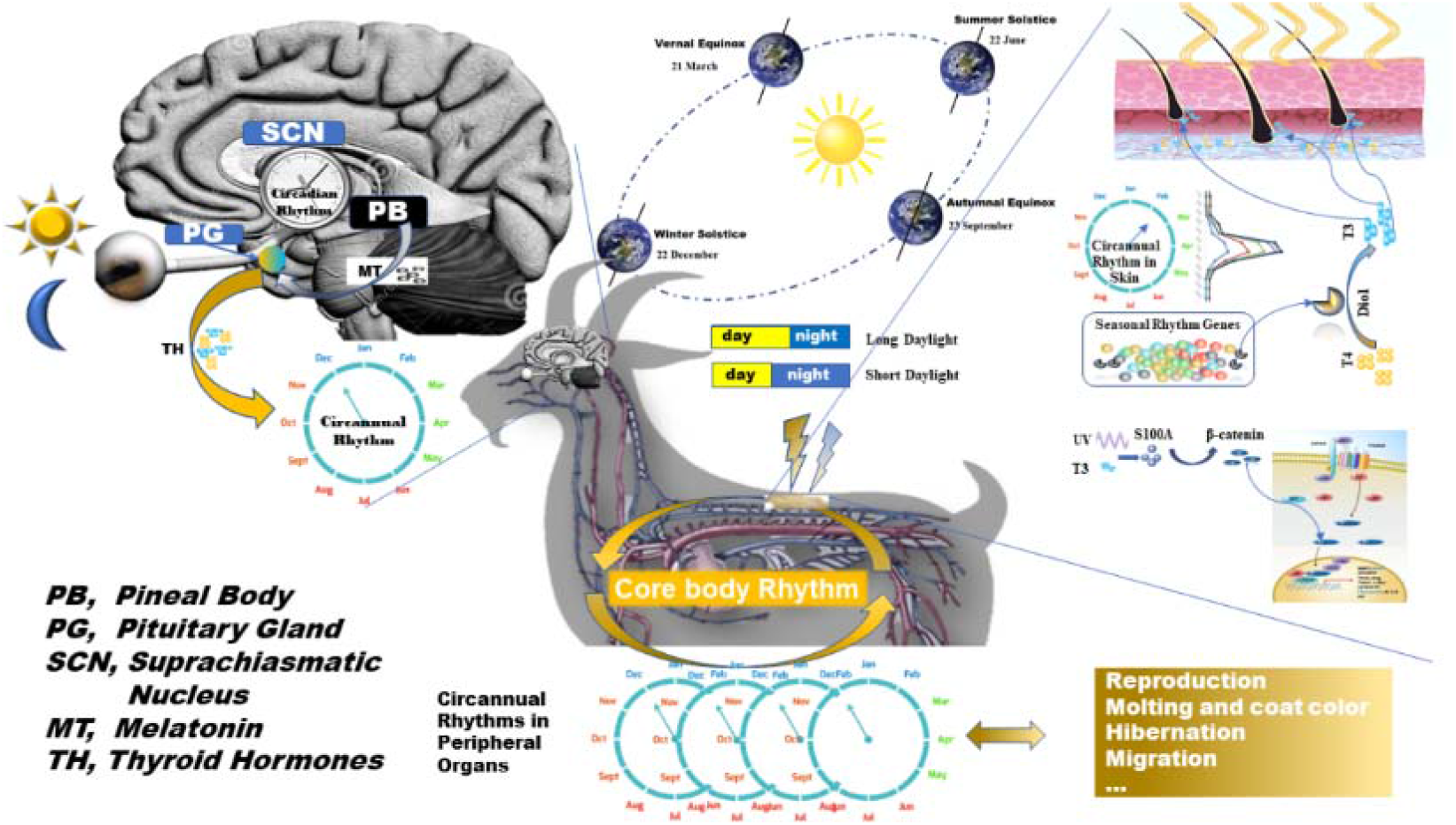
The Hypothesis of the Skin Circannual Rhythm Interaction with the Photoperiod.

Taking together, we provided a hypothesis to describe how goat skin makes the own rhythm and gets the clue from the environment factor. The melatonin, produced by the pineal body, is dependent upon the transmission of the light through the eye that is decided the circadian rhythm. Melatonin could inhibit the activity of the thyroid gland through the pituitary gland and decrease the secretion of thyroid hormone. Then the content of thyroid hormone in blood circulation maintains the core body circannual rhythm of a goat. When the compounds of TH, T3 and T4, reach to the peripheral organs such as skin, the located deiodinase regulating the transformation from the T4 to T3. The expression level of deiodinase or other molecules are affected by the environment factor such as the photoperiod. Finally, the interaction of transcription factor in SRGs and miRNAs regulated the gene expression profile and physiological process in the skin to adapt to the season change. However, it needs more advanced functional experiments in the future to identify the details of how the skin gene expression profile adapts to season variation.

## Method and Materials

### Experimental animals and the location

Cashmere goat is a famous livestock species for the cashmere. We collected skin samples of body side from three Inner Mongolia Cashmere Goats. As the experimental goats were located at Tumed Zuoqi, Huhhot, China (N40°30′22.89″E110°54′47.29). We collected skin samples up to ~1.5 cm^2^ from body side of cashmere goat. Then the samples were immediately stored at liquid nitrogen before RNA extraction. Skin samples were collected at noon on the first three days of every month for one year, from December 2014 to December 2015, from the bodyside of three natural goats. In total, transcriptomes of 36 skin samples were obtained by Illumina sequencing (RNA-Seq).

### SMRT sequencing of full-length transcriptome of skin

We collected body side skin and ear skin from three Inner Mongolia Cashmere Goats at four time points (Jan, Apr, Jul, Oct). The RNA samples were extracted with Trizol from skin sample. Equal amounts of total RNA from each skin sample were pooled prior to library construction. The SMART sequencing library was constructed according to the Iso-Seq protocol. Full-length cDNAs of mRNAs were synthesized by SMARTer™ PCR cDNA Synthesis Kit. PacBio Iso-Seq libraries were prepared following the PacBio Standard Library prep protocol. Sequencing was performed on a PacBio RsII instrument. Both PacBio sequencing, Illumina sequencing and sRNA was performed by Biomarker Technology Co. (Beijing, China).

Raw reads were processed into error corrected reads of insert (ROIs) using Iso-seq pipeline with minFullPass=3 and minPredictedAccuracy=0.9. Next, full-length, non-chemiric (FLNC) transcripts were determined by searching for the polyA tail signal and the 5’ and 3’ cDNA primers in ROIs. The high-quality RNA-seq reads were used to further correct FLNC CCSs by proovread 2.13 using the default parameters[42]. ICE (Iterative Clustering for Error Correction) was used to obtain consensus isoforms. FL consensus sequences from ICE was polished using Quiver. High quality FL transcripts were classified with the criteria post-correction accuracy above 99%. FL consensus sequences were mapped to reference genome using GMAP. Mapped reads were further collapsed by pbtranscript-ToFU package with min-coverage=85% and min-identity=90%. 5’ difference was not considered when collapsing redundant transcripts.

### Annual Transcriptome of Cashmere Goat Skin

Total RNA was extracted using a RNeasy Mini Kit (Qiagen) from a skin sample. The amount of RNA was measured using a Nanodrop or Qubit 2.0 (Thermo Fisher Scientific), and the quality was assessed using an Aglient 2100 (Agilent Technologies). The RNA-seq library was constructed with 150bp paired-end sequences for each sample, according to the standard protocol provided by Illumina, Inc. (San Diego, CA, USA). FastQC was used to calculate the quality control statistics for the data generated by the Illumina HiSeq4000, and the resulting libraries were then subjected to paired-end sequencing (2 × 150bp).

To calculate the expression level of skin genes, Salmon [13] software was utilized based on the whole-length transcripts of goat skin as a reference sequence. Then, differential expression genes(DEGs) were detected by EdgeR package of R [43]. We normalized the expression matrix of DEGs by dividing each value by the gene of its maximum value observed in any sample for that gene. To analyze the annual transcriptomes of goat skin, we used the MetaCycle [14], an integrated R package, to evaluate periodicity in large scale data, to find the annual rhythm expressed genes.

### Seasonal Skin sRNA sequencing

We collected 8 body side skin samples from three Inner Mongolia Cashmere Goats at eight-month (Jan, Apr, May, Jul, Aug, Sep, Oct, and Dec) and classified samples as winter (Dec and Jan), spring (Apr and May), summer (Jul and Aug), fall (Sep and Oct). A total amount of 2.5 ng RNA per sample was used as input material for the RNA sample preparations. Sequencing libraries were generated using NEBNext^®^ small RNA Sample Library Prep Kit for Illumina (NEB, USA) following the manufacturer’s recommendations and index codes were added to attribute sequences to each sample. After cluster generation, the library preparations were sequenced on an Illumina Hiseq 2500 platform and paired-end reads were generated. miRDeep2 was utilized to detect mature miRNAs and predict novel miRNAs [16]. The differential expressed miRNAs (DEMs) were identified by the DESeq package in the R language [44].

### Bio-information Analyses

To analyze the role of transcript factor in SRGs, we utilized the AnimalTFDB to investigate the regulator motifs in 5’FTR of SRGs. ClueGO, a plugin of the Cytoscape platform [45], was used to classify the function of seasonal rhythm genes. Enrichment/Depletion (Two-sided hypergeometric test), Correction Method (Bonferroni step down), Min GO Level (3), Max GO Level (12), GO Fusion (false), GO Group (true) were set in ClueGO. MetaCycle[14] was utilized To analyze the rhythm of DEGs of goat skin. We tried from 4 months to one year as a length of interesting rhythms. And then to bridge the relationship between DEMs and SRGs, miRanda [46] was used to find the binding site in SRGs sequence for DEMs. Then, interaction networks of DEMs and SRGs was drawn using the Cytoscape.

## Declarations

### Ethical Approval and Consent to participate

My manuscript report data collected from cashmere goat skin. The sampling procedure was following the standards of the Animal Care and Use Committee in Inner Mongolia University for Nationalities, China.

### Consent for publication

All authors provided consent for publication.

### Availability of supporting data

The multiple Omics data used; analysis pipeline used in this study will be made available from the corresponding author upon request.

## Authors’ contributions

Conceived and designed the experiments: JW. Performed the experiments: JW YL HS. Analyzed the data: HG JW LD. Contributed reagents/materials/ analysis tools: YL DW CL. Wrote the paper: JW.

## Authors’ information

Jianghong Wu (, College of Animal Science and Technology, Inner Mongolia University for Nationalities, Tongliao, 028000, China; Inner Mongolia Academy of Agricultural & Animal Husbandry Sciences, Hohhot, 010031, China)

Yin li (, College of Animal Science and Technology, Inner Mongolia University for Nationalities, Tongliao, 028000, China)

Husile Gong (, College of Life Science, Inner Mongolia University for Nationalities, Tongliao, 028000, China)

Dubala Wu (, College of Animal Science and Technology, Inner Mongolia University for Nationalities, Tongliao, 028000, China)

Chun Li (, College of Animal Science and Technology, Inner Mongolia University for Nationalities, Tongliao, 028000, China)

Bin Liu (, Inner Mongolia Academy of Agricultural & Animal Husbandry Sciences, Hohhot, 010031, China)

Lizhong Ding (, David Geffen School of Medicine, University of California, Los Angeles, CA 90095, USA)

---

**Supplement file 1.**
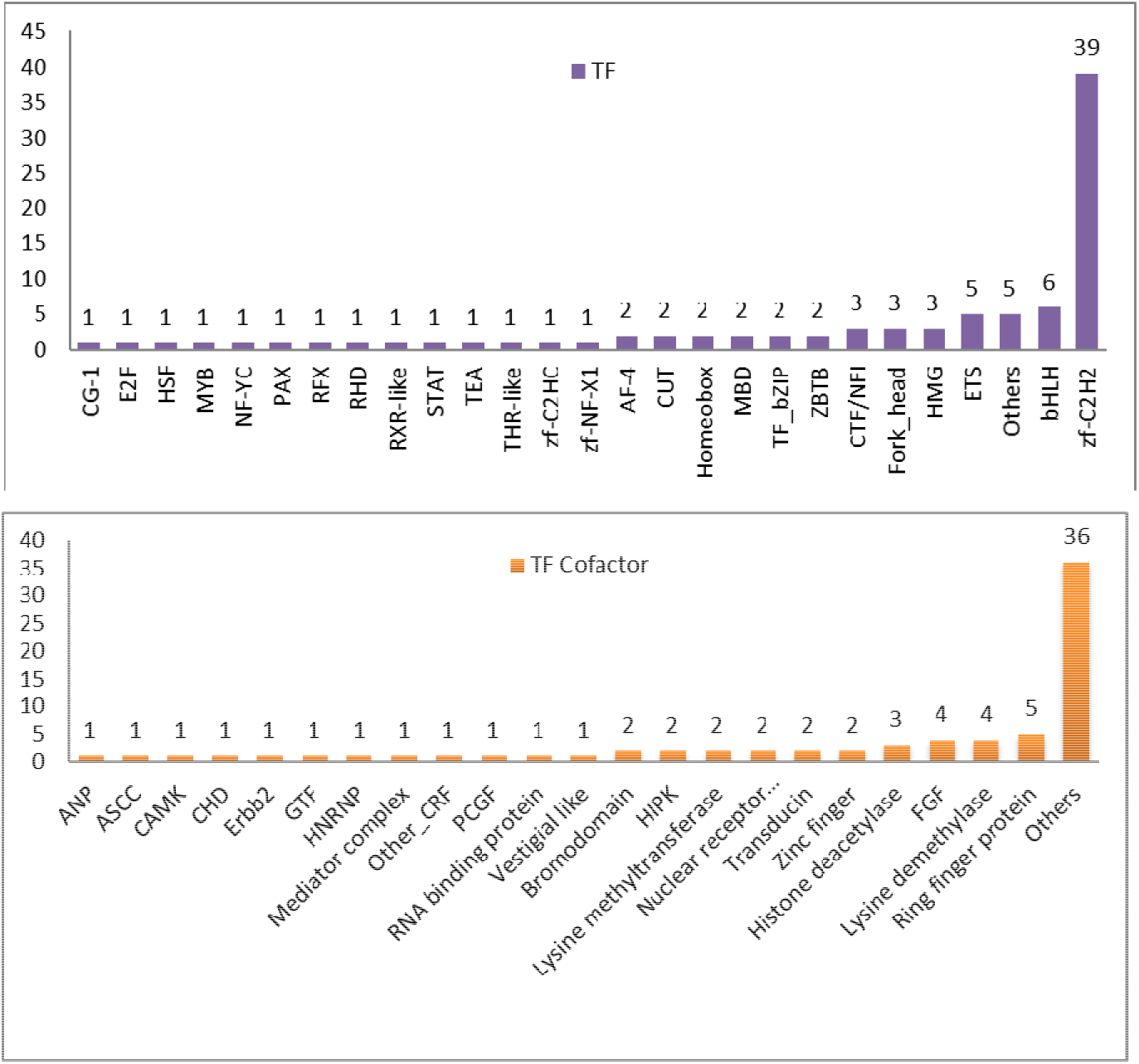
the class of transcript factor (top) and TF cofactor(bottom) for SRGs

**Supplement file 2.**
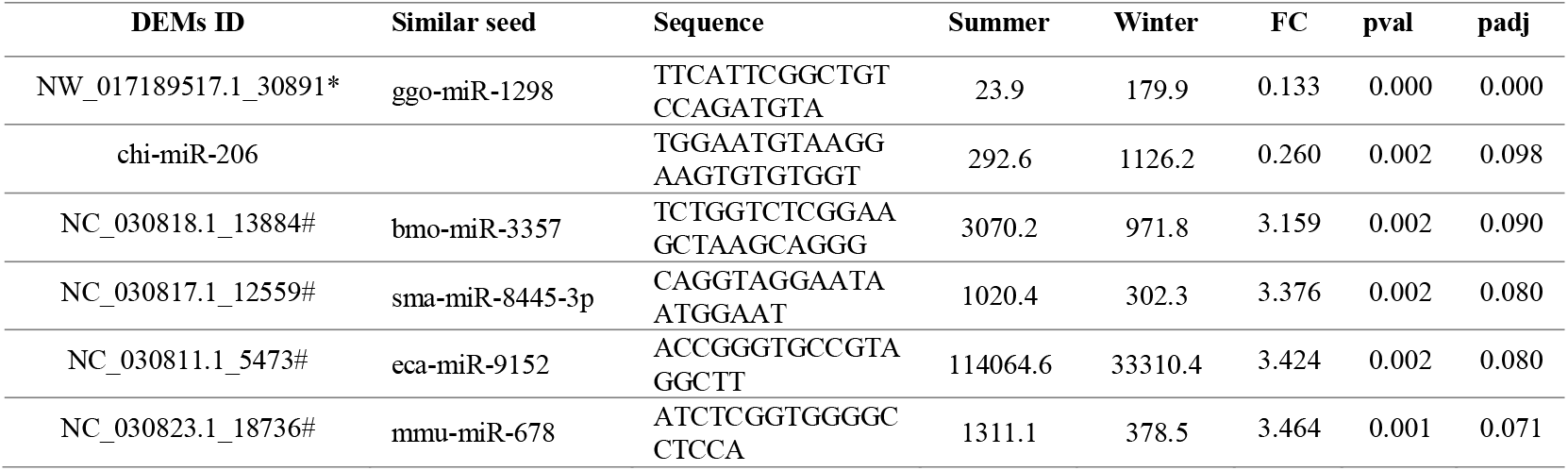
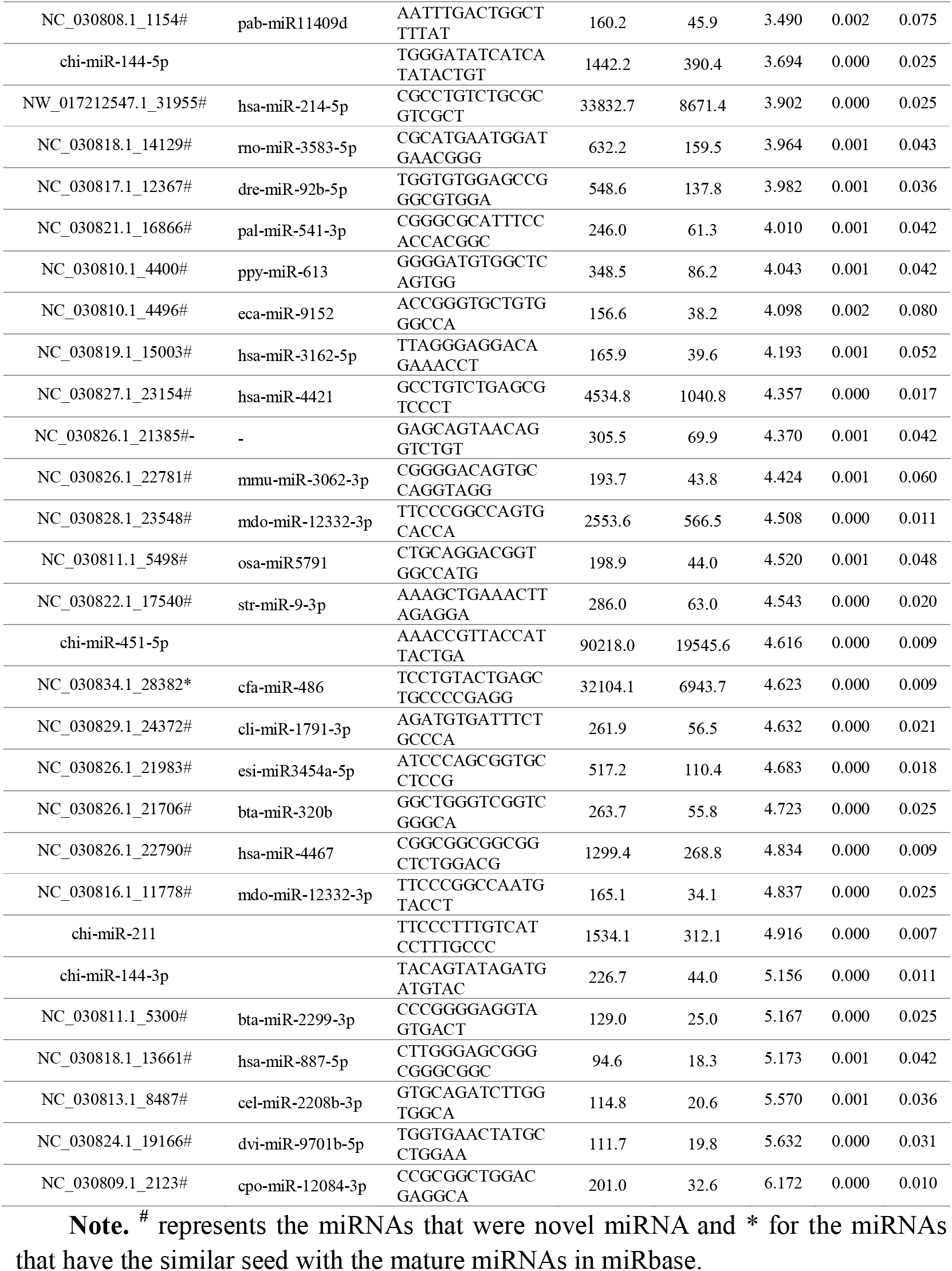
the information of differential expressed miRNAs between summer and winter

